# Linking drug target and pathway activation for effective therapy using multi-task learning

**DOI:** 10.1101/225573

**Authors:** Mi Yang, Jaak Simm, Chi Chung Lam, Pooya Zakeri, Gerard J.P. van Westen, Yves Moreau, Julio Saez-Rodriguez

## Abstract

Despite the abundance of large-scale molecular and drug-response data, the insights gained about the mechanisms underlying treatment efficacy in cancer has been in general limited. Machine learning algorithms applied to those datasets most often are used to provide predictions without interpretation, or reveal single drug-gene association and fail to derive robust insights. We propose to use Macau, a bayesian multitask multi-relational algorithm to generalize from individual drugs and genes and explore the interactions between the drug targets and signaling pathways’ activation. A typical insight would be: “Activation of pathway Y will confer sensitivity to any drug targeting protein X”. We applied our methodology to the Genomics of Drug Sensitivity in Cancer (GDSC) screening, using gene expression of 990 cancer cell lines, activity scores of 11 signaling pathways derived from the tool PROGENy as cell line input and 228 nominal targets for 265 drugs as drug input. These interactions can guide a tissue-specific combination treatment strategy, for example suggesting to modulate a certain pathway to maximize the drug response for a given tissue. We confirmed in literature drug combination strategies derived from our result for brain, skin and stomach tissues. Such an analysis of interactions across tissues might help target discovery, drug repurposing and patient stratification strategies.

## Introduction

Translating preclinical models into actionable insight is essential for more personalized treatments. Despite the wealth of omics data since a few decades, our ability to decipher the mechanisms underlying drug response has been much less effective^1^. This is particularly apparent in cancer. Large scale drug screenings have been a major resource for drug discovery. In particular, public drug screening projects such as the Genomics of Drug Sensitivity in Cancer (GDSC)^2^, the Cancer Therapeutics Response Portal (CTRPv2)^3^ and the Cancer Cell Line Encyclopedia (CCLE)^4^ have generated drug response data for hundreds of drugs and around one thousand cell lines. The main objective of these datasets is to shed light on the molecular mechanisms regulating drug response.

From this data, machine learning is widely used to predict drug response on the treated cell lines. Most of the analyses consist of building a model for one drug at a time, which has limited power given the relative low number of samples. If we can bring together all drugs in a single model, we can learn common patterns reflecting the underlying mechanisms. Towards this end, multitask type algorithms which use information gained in one task for another task are a promising approach, that have been recently applied to drug sensitivity prediction from large scale drug screenings^5,6,7^. Methods ranging from standard random forest^6^ to Kernelized bayesian matrix factorization^8^ and trace norm multitask learning^5^ have been used to predict drug response by integrating genomic features for cell lines, as well as target and chemical information for drugs. While many multitask algorithms perform better than standard methods, interpretability is often challenging.

The motivation of our work was to leverage the power of multitask learning to provide novel insights into the molecular underpinnings of drug response. Towards this end, we applied a multitask learning strategy for drug response prediction and feature interaction, using the tool Macau^9^. Our algorithm tries to learn multiple tasks (predicting multiple drugs) simultaneously and uncovers the common (latent) features that can benefit each individual learning task^10^. We focused on gene expression as molecular input data, using it to estimate activities of signaling pathways, along genetic aberrations. For the drugs, we chose their nominal target as the key feature. We applied our methodology to the Genomics of Drug Sensitivity in Cancer^11^ (GDSC) cell line panel with drug response (IC50) of 265 drugs on 990 cell lines. The interactions between protein targets and signalling pathways’ activities support a personalized treatment strategy to, for example, determine how to modulate a certain pathway to maximize the drug response. To portray tissue specificity in cancer treatments, we explored the differences of interactions across tissues with different compounds. Analyzing those interactions across many tissues can enable patient stratification, drug repositioning, and drug combination selection.

## Results

### Drug response prediction in different settings

While our aim was to use Macau to obtain interpretable results rather than improve predictability, we first compared the performance of Macau to standard linear regression (ridge/lasso) and tree based non linear regression such as Random Forest and XGBOOST (**Supplementary Table S2**). When building models to predict drug response taking into account multiple drugs and cell lines, one can define four differents settings which mirror different use cases (**Supplementary Fig. S1**). As cell line descriptor, we used gene expression, as well as PROGENy^12^ pathway scores. PROGENy is a data driven pathway method aiming at summarizing high dimensional transcriptomics data into a small set of pathway activities (**Methods**). The 11 PROGENy pathways currently available are EGFR, NFkB, TGFb, MAPK, p53, TNFa, PI3K, VEGF, Hypoxia, Trail and JAK STAT. For setting 1, 2 and 4, we used 10 fold cross validation, repeated 10 times. We define the prediction performance as the Pearson’s Correlation (r) of observed versus predicted drug response (IC50). We also provide the comparison results for gene expression, SNP/CNV and ECFP4 fingerprint as input in the **Supplementary analysis 1**.

#### Setting 1

Prediction of new cell lines for existing drugs

The meaning of this framework is to start with a subset of drugs and assign them to the right patient, e.g. new patients based on their genomic information (**Supplementary Fig. S1a, Supplementary Table S1**). There is no significant difference between Macau (r=0.30), Ridge (r=0.30), Random Forest (r=0.31) and XGBOOST (r=0.31), see **Supplementary Table S2**.

#### Setting 2

Prediction of new drugs on existing cell lines

A second important scenario is to predict the effect of a new drug on a set of patients based on the side information of the drug. If the new drug is predicted to be better than the existing ones, then a therapeutical switch can be considered. The concept of “new drug” is relative to the patient, it can concern existing drugs which have never been used for a patient group (**Supplementary Fig. S1b, Supplementary Table S1**). By using drug target as input, Macau outperforms Ridge, Random Forest and XGBOOST with a performance of 0.42 against 0.12, 0.38 and 0.19, respectively, all p values < 2.2e-16 (**Supplementary Table S2**). Of note, Random Forest performed significantly better than Ridge and XGBOOST with r=0.38 against r=0.12 and r=0.19, respectively, all p values < 2.2e-16.

#### Setting 3

Prediction of existing drugs and existing cell lines (**Supplementary Fig. S1c, Supplementary Table S1**)

In this setting we solve an imputation problem, where the test set is randomly chosen from the drug response matrix. We can use side information from both sides to improve the result. We tested setting 3 on GDSC (**Supplementary Table S3**) datasets. In overall, we were able to get an excellent prediction: mean r=0.932 with 90% of the data as training set and 10% and even r=0.834 with 99% as test set.

#### Setting 4

Prediction of new drugs on new cell lines (**Supplementary Fig. S1d, Supplementary Table S1**)

This setting aims at predicting a new drug’s effect on a new cell line solely based on drug target information and whole transcriptomics, hence a very challenging task. In this analysis, all algorithms were used in multitask mode. Macau performed at r=0.44 (sd=0.15), not significantly different from Elastic net (r=0.41, sd=0.12, p value=0.24) and Random Forest (r=0.42, sd=0.17, p value=0.34)(**Supplementary Table S2**). Macau outperforms XGBOOST (r=0.39, sd=0.17), with respective p values of 0.0067.

We also performed a benchmark in a tissue-specific setup. We compared for 16 tissues, the prediction using GEX, PROGENy and SNP/CNV (**Supplementary Table S4**). For most of the tissues, the pearson correlation of observed versus predicted IC50 is close to 0.4. Gene expression does not perform significantly better than PROGENy in most tissues, except for colon (p-value = 0.02), liver (p-value = 0.01), soft tissue (p-value = 0.03) and stomach (p-value = 0.002). Compared to SNP and CNV, gene expression performs significantly better for 14 out of the 16 tissues.

In summary, our multitask learning achieves a similar or better predictability performance than standard methods (single task and multitask) across all settings. We can now focus on the insights generated by the Macau model.

### Exploring underlying interactions with Macau

Macau is a Scalable Bayesian multi-task learning algorithm which can incorporate millions of features and hundred millions of observations^9^. In traditional machine learning analysis, we predict the response variable based on descriptive features of the samples. For instance, in drug screening experiments where cell lines are treated by drugs, the effect of a certain drug X is predicted from the mRNA expression of a gene Y via regression. With Macau, we unveil the interaction matrix of the drugs’ feature (for example, protein target) with the cell lines’ feature (e.g. transcriptomics, pathway activity). A typical insight would be: “Upregulation of gene Y correlates with drug sensitivity when targeting protein X” (**Figure 1a**, **Methods**). If this were a causal relationship, it could mean that upregulation of gene Y confers sensitivity to any drug targeting protein X. We refer to this from now on as feature interaction analysis. Such analysis gives hints about the drug’s mode of action, by uncovering how acting on one protein affect the drug response and in which conditions (gene/pathway status).

**Figure 1.**
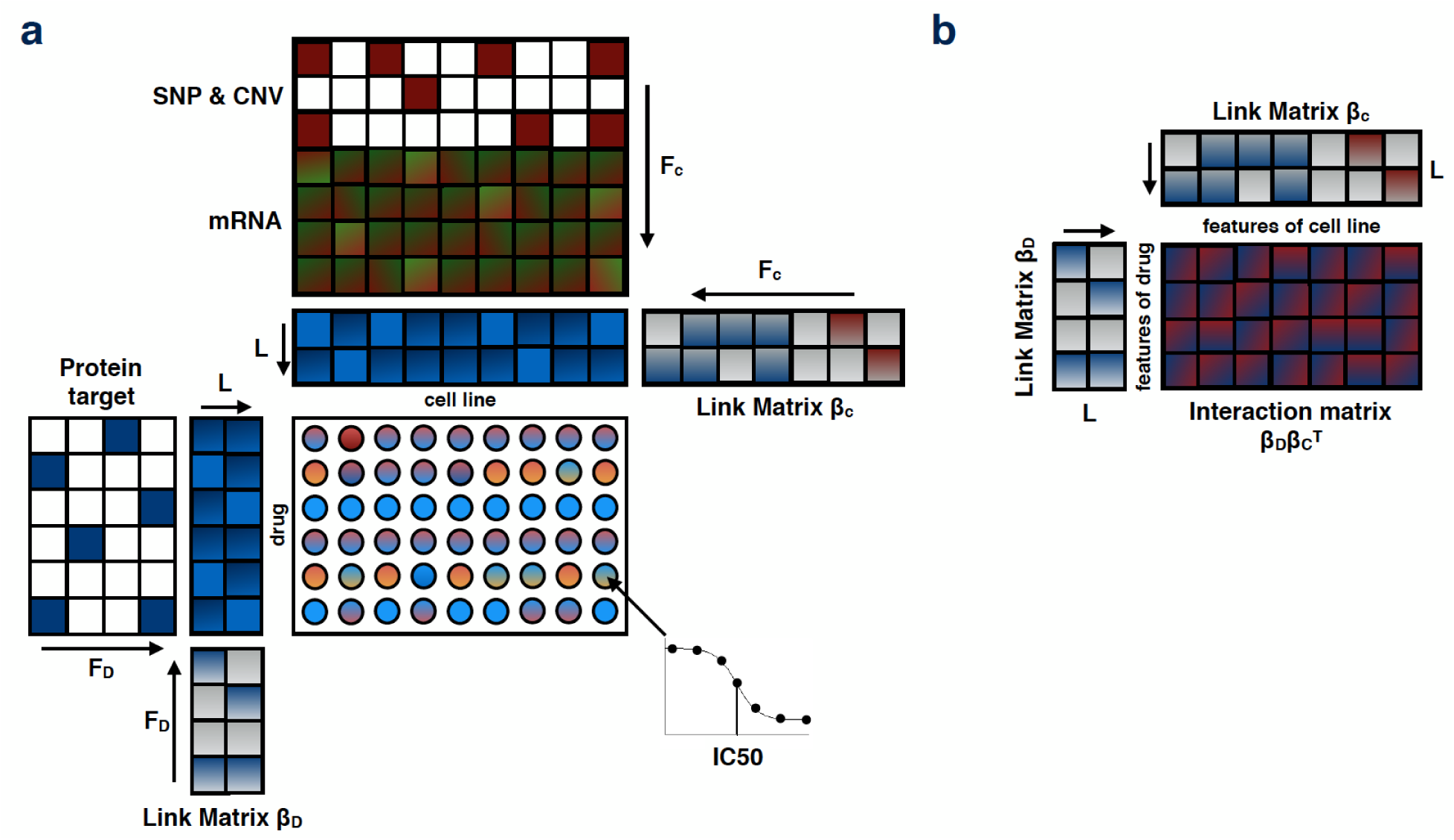
Macau factorization model: **(a)** The drug response (IC50) is computed by 2 latent matrices. Each of them is being sampled by a Gibbs sampler. In presence of additional information (side information), the latent matrix is predicted by a multiplication of a link matrix and the side information matrix. Arrows in this figure indicate the matrix multiplication. **(b)** By multiplying the 2 link matrices, we obtain the interaction matrix, which is the interaction between the features of the drugs with the features of the cell lines.

In our feature interaction analysis, we used manually curated protein targets for the drug side obtained from the GDSC website (https://www.cancerrxgene.org/downloads). For the cell line side, we used PROGENy scores. We computed with Macau the interaction matrix with features of the drugs on the rows and features of cell lines on the column (**Methods**).

### Feature interactions: Tissue specific analysis

Using features on both sides of the drug response matrix, we can measure the interaction between features of drugs and features of cell lines, by taking into account all drugs and all cell lines in a generalized model. Based on the results of Macau’s performance in drug response prediction in different settings (**Supplementary Fig. S1, Supplementary Fig. S2**), we chose to use PROGENy pathway activity for cell lines due to performance and interpretability reasons, and protein targets for drug side. We confirmed our choice for PROGENy by demonstrating it’s superior predictive performance compared to other pathway methods (**Methods, Supplementary Fig. S9**), in agreement with its superior ability to find statistical associations with drug response^12^.

We performed a feature interaction analysis of drug target - PROGENy pathways for all 16 tissues (**Supplementary Fig. S3**) and assessed the significance of the interaction matrices (**Methods**). We then examined the interactions to derive biological insights in terms of biomarkers, potential drug combination, and drug repositioning. We will highlight in the following examples with the support from literature for four tissues (bone, brain, skin and stomach; **Figure 2**), and for four additional tissues (aerodigestive tract, breast, SCLC lung and pancreas) we provide the results in the **Supplementary analysis 2**.

**Figure 2.**
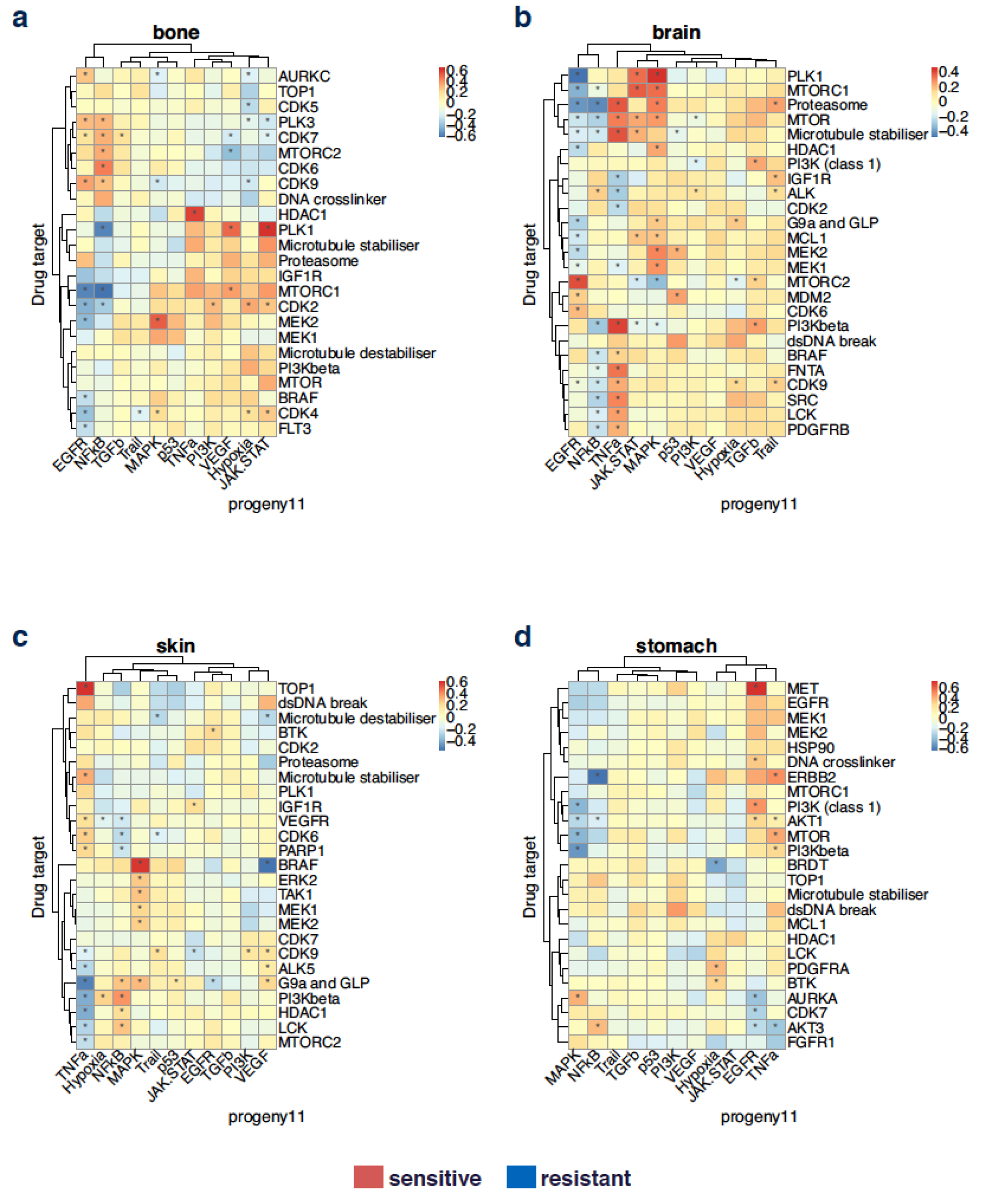
Tissue specific analysis of interaction matrix. We used target on drug side and pathway activity on cell line side and analyzed all tissues in the GDSC panel with at least 20 samples, and display the targets which have an interaction for at least 1 pathway in the top 5% absolute value. We subset the targets a second time by keeping the top 25 targets with the highest variance across the pathways in term of interactions. Here, we highlight 4 representative tissues: **(a)** Bone. **(b)** Brain. **(c)** Skin. **(d)** Stomach.

#### Bone (Figure 2a)

From the heatmap results, we observe that cells are sensitive to MEK1/MEK2 inhibition when MAPK pathway is activated. This is an expected result based on the general paradigm of oncogenic addition, and indeed it has been reported that MEK inhibition induces apoptosis in osteosarcoma cells with constitutive ERK1/2 phosphorylation^13^.

#### Brain (Figure 2b)

i. In our results, EGFR pathway activation correlates with drug sensitivity when MTORC2 is targeted. This agrees with the fact that EGFR activates mTORC2-NF-κB pathway in glioblastoma cells promoting growth, making these cells likely addicted to MTORC2^14^.
ii. Cells with active EGFR signaling are resistant to PLK1 inhibitors. Therefore, one could hypothesize that blocking EGFR pathway while targeting PLK1 could lead to synergistic effect. In agreement with that, PLK1 and EGFR inhibitor have been described as orthogonal therapeutic agents in glioblastoma, with enhanced tumoricidal activity when combined^14,15^.

#### Skin (Figure 2c)

i. Activation of TNFa pathway correlates with drug sensitivity when TOP1 is targeted. The mechanism underlying this relationship is less clear, but repression of TOP1 activity inhibited IFN-β- and TNFa-induced gene expression, suggesting a link between these processes^16^.
ii. We observe in skin that MAPK activation correlates with drug sensitivity when targeting BRAF. This is expected as BRAF activates the MAPK pathway, and the association is only found in skin, where BRAF inhibitors have proven successful in the clinic, specifically by blocking the mutant form BRAF^V600E^^17^. Conversely, activation of VEGF pathway confers resistance when targeting BRAF. This suggests that blocking VEGF can have a synergistic effect with targeting BRAF, and in fact Dual BRAF^V600E^ and VEGF targeting has been shown to provide a combinatorial benefit against BRAF^V600E^ mutants tumor growth *in vivo^18^*. This case illustrates the use of our approach to identify drug combinations by targeting a different pathway.

#### Stomach (Figure 2d)

One striking example is increased sensitivity by targeting MET when the EGFR pathway is activated. In agreement with our result, combination of anti MET/EGFR has been proven efficient in MET-amplified gastroesophageal xenopatient cohort^19^.

We then seek literature support for the interaction matrix in an automatic and unbiased way. For each tissue, we searched for the number of publications containing the keywords: “target” AND “pathway” AND “tissue of interest”. For an interaction matrix of 102 drug target and 11 PROGENy pathways, we obtained a Pubmed matrix of the same dimension. We then took the absolute value of the interaction weight. We removed triplets with no publication found, triplets with q-value of the interaction weight of 1 and triplets with similar name for protein and pathway. We then correlate the number of publications against the interaction weight (**Supplementary Table S8**). For 15 of the 16 tissues, the correlation is positive. For 10 tissues, the false discovery adjusted p value < 25%. Hence, the number of publication tend to increase with the absolute value of the interaction weight.

In summary, we could find literature support for the results of the tissue specific analysis, suggesting that insights generated from feature interaction analysis could have clinical impact. We also confirmed several drug combination strategies (targeting the same pathway or different pathways) and we argue that the target - pathway interaction heatmap can be powerful tool for deriving combination strategies simply by it’s visualization power. We will now focus on cross tissue exploration of the result.

### Therapeutic applications of the interactions

#### Deriving biomarker and drug combination strategy

If the interaction between a pathway activity and the drug efficacy is causal and not just a correlation, modulating the pathway would affect the drugs’ effect. For each tissue, we selected two target-pathway pairs, one sensitive association and one resistant association. We plotted the IC50 against the corresponding pathway for a drug targeting the corresponding protein target (**Supplementary Fig. S4**). In 50% of the cases, the correlation between the IC50 and the PROGENy score were significant (p<0.05). The pathway activity could potentially be used as biomarker of drug response for a given tissue.

Another use case would be drug combination: if a pathway activity correlates with resistance to a given drug, targeting the pathway might increase the efficacy of the drug, as some of the examples shown in the previous section. As an additional example, in lymphoma, activity of the NFkB pathway, which is constitutively deregulated in lymphoma development^20^, correlates with resistance when using antimetabolites, a common type of chemotherapy (**Figure 3, Supplementary Fig. S3k**). Thus, blocking NFkB may restore sensitivity to antimetabolite drugs. Interestingly, in Non Hodgkin lymphoma, antimetabolites are used together with corticosteroids (protocol CVAD + Methotrexate and Cytarabine). As corticosteroids inhibit NFkB^21^, this could explain the combination.

**Figure 3.**
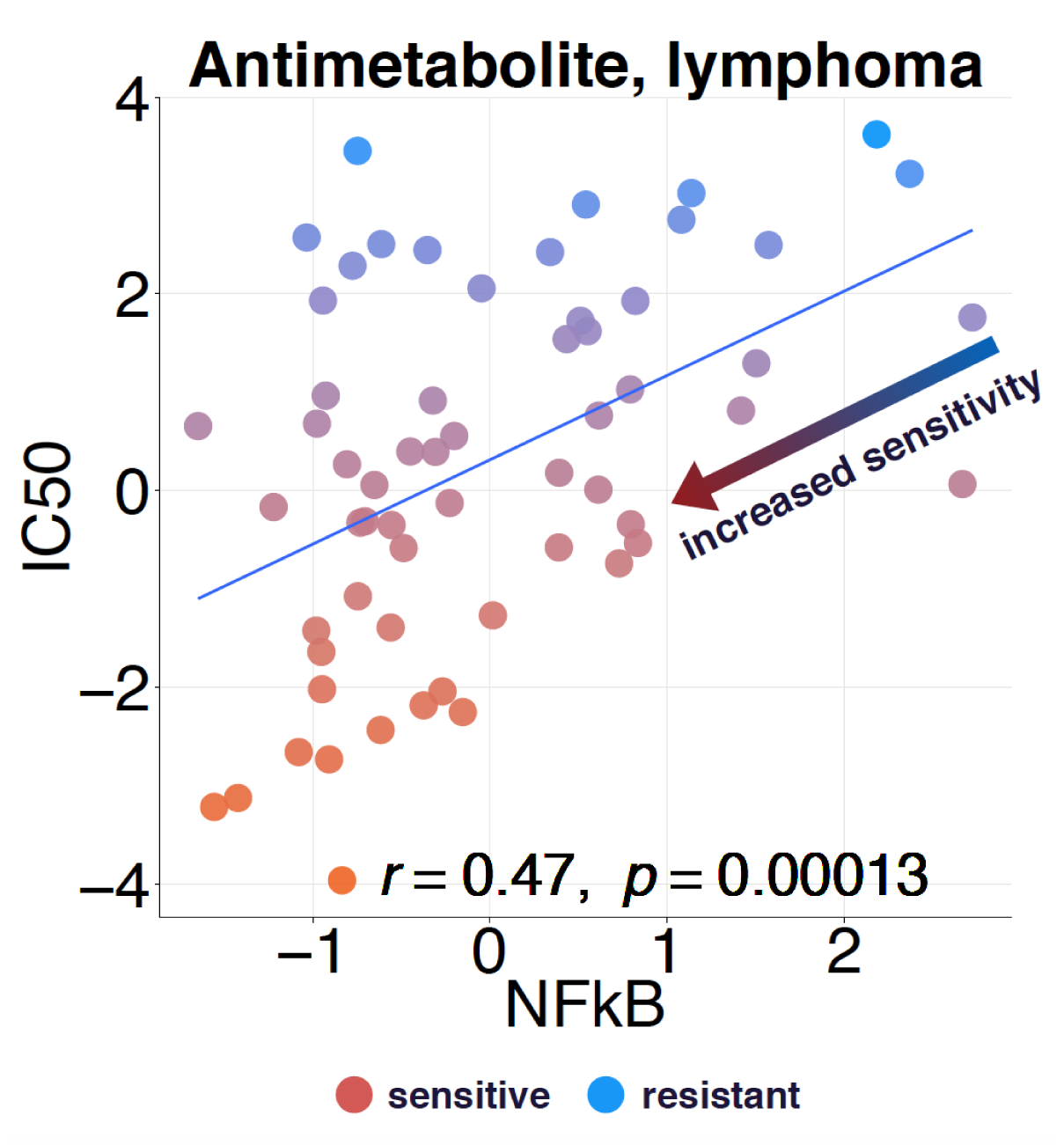
Increasing sensitivity and overcoming resistance. From tissue specific interaction matrix of lymphoma, we chose the top hits Antimetabolite - NFkB (as target - pathway pairs). We plot the IC50 of drug Cytarabine against NFkB pathway’s activity.

#### Harnessing tissue variability of interactions

In order to explore dissimilarities across tissues, for each pathway-target pair, we selected the tissue where it has the highest interaction weight and the tissue with the lowest weight. Then, we kept the pairs with smallest difference in absolute value between maximal and minimal weight. The objective is to find target-pathway pairs which have the greatest and most antagonistic effect for two different tissues (**Supplementary Table S6**). For instance, NFkB confers high sensitivity in breast but resistance in stomach to drugs targeting ERBB2 (**Supplementary Fig. S5**). In most cases we could discern an antagonistic behavior from one tissue to the other, except for EGFR-DNA damage pairs.

We next explored the similarities between tissues. We started with a matrix of dimension 16 tissues x 1122 pathway-target pairs, and then subset the interactions by taking only into account the pathway-target pairs for which at least one tissue appears in the top 5% absolute value. Finally, we rank the remaining pairs by the variance of their interactions across the 16 tissues and keep the lowest 30. In this highest interaction heatmap (**Figure 4a**), we highlight the pathway-target pairs which confer drug sensitivity for many tissues. This allows the use of the same drug in the same condition, but on a different tissue. To find the dissimilarities between tissues, we followed the previous steps, but instead keeping the top 30 pairs with the highest variance of interactions across tissue. This divergent interaction heatmap (**Figure 4b**) displays the pathway-target pairs which have a huge variance across tissues.

**Figure 4.**
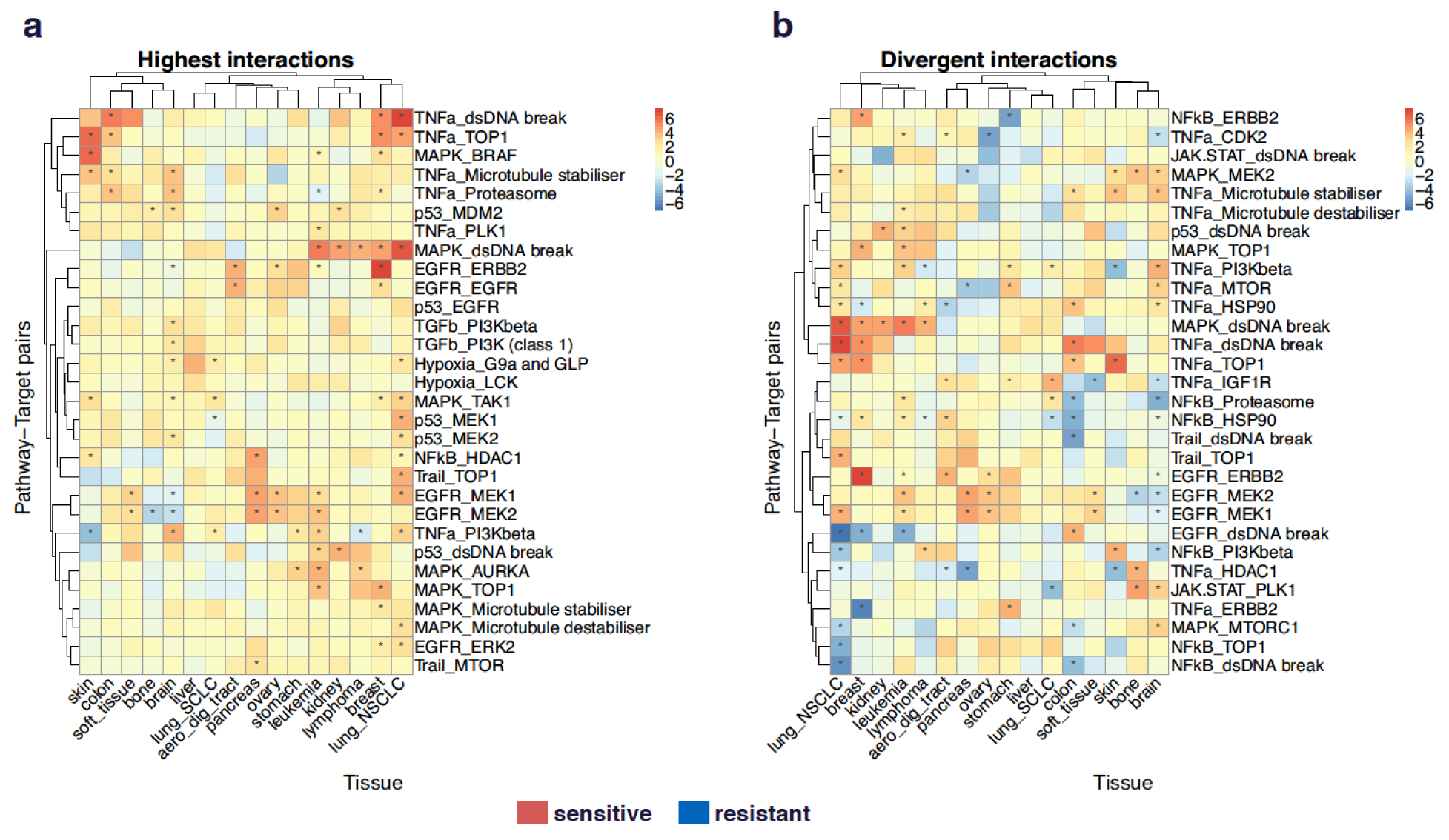
Feature interaction analysis across tissues. **(a)** Highest interactions. We vectorize all cancer specific interaction matrices between target and PROGENy pathways and obtain a matrix of dimension (number of tissues x number of pathway target pairs). We do a first subsetting by taking only into account the pairs for which at least one pathway appears in the top 5% absolute value. We then keep the 30 pathway-target pairs with the highest mean value across tissues in term of interaction. **(b)** Divergent interaction. Same as previously, except that we keep the top 30 pairs with highest variance across tissues.

### Validation on external datasets

In order to test the robustness of our findings, we applied our method to CTRPv2^3^ dataset (481 compounds x 860 cell lines) as its size (number of drugs and cell lines) is comparable to GDSC. For each tissue, we compared for GDSC and CTRPv2 the interaction matrices between drug target and PROGENy pathways. We considered 39 protein targets which are in common and targeted by at least two drugs in both datasets. For aerodigestive tract tissue, the 71 points in the interaction matrix (out of 401) which have a q-value < 1 (Benjamini-Hochberg Yekutieli; see **Methods**) in both GDSC and CTRP datasets have a pearson correlation of 0.55 (p=6.9e-07) between studies. If we set the threshold of q-value at 0.40, the correlation would be 0.68 (p=0.01004) but with 13 data points, and only 4 with q-val < 0.25. Similarly, we obtained for the other tissues in common between both datasets (q-val<0.40): 0.82 (p=3.8e-07, breast), 0.34 (p=0.41, colon), 0.81 (p=0.19, ovary), 0.61 (p=0.06, pancreas), 0.63 (p=0.022, skin) and 0.33 (p=0.31, stomach). In overall, the two datasets are in agreement for the common target-pathway interactions.

### Interaction models from predicted drug targets

Drugs are known for binding to multiple proteins, but our interaction models are based on a literature curated target list, where each drug binding to a handful of proteins. In order to assess the full spectrum of potential protein targets, we used deep learning to predict drug protein binding from the ChEMBL database^22^ (**Supplementary Method**). We applied this method to the drugs used in the GDSC dataset and predicted drug protein binding for 696 proteins targets. We then ran the target - pathway interaction analysis (**Supplementary Fig. S8**). For bone cancer, we found that targeting protein NTSR1 is associated with drug sensitivity when the Hypoxia pathway is activated (**Supplementary Fig. S8c**). Protein Neurotensin Receptor 1 has been proposed as a potential therapeutic target^23^. The fact that a protein which does not belong to the manually curated list can be a hit, suggests that those analysis could be used as resources for drug target discovery.

## Discussion

In this paper, we provide a powerful machine learning framework for large scale drug screenings to find interactions between the drugs’ and the cell lines’ characteristics. We focused on exploring how pathway activities modulate response to drugs targeting specific proteins.

In traditional analyses, findings are typically about the association between a drug and a gene. Such approach has the limitations that a gene alone may not capture the entire complexity of the signaling landscape, and the drug may not be very relevant and not used after the publication, therefore the insight is lost and more generalizable insights are desirable.

To overcome these issues, we introduced the feature interaction analysis in cancer specific settings. We rely on a data driven pathway method (using perturbation experiments) that has proven to be efficient at estimating pathway status^12^ from gene expression. We explored the tissue specificity of target - pathway pairs, and we found literature support for many of our findings in term of the effect of targeting a specific protein in presence of a pathway’s activation for a certain cancer type. This would not have been possible without an efficient way to reduce high dimensional omics data into a small and interpretable subset of pathways. Our results show how multitask learning can handle large scale experiments and derive interpretable insights that may ultimately improve clinical decision and therapeutical choices.

There are several limitations to this study: First, the quality of the insights depends on the quality of the target pathway interaction. The performance (in setting 4) is ~0.4 for breast and colon cell lines, and up to 0.45 for skin and aerodigestive tract (**Supplementary Table S4**), which is still far from perfect. A significant part of mechanism are not explained by those pathways. We could address this issue by, for example, expanding the PROGENy pathways. Second, one limitation of the GDSC panel for our analysis is that it adjusts the drug concentration range for each compound individually, to have a few cell lines responding, while the large bulk of cell lines does not respond, which makes the drug sensitivity in cell lines a relative concept. Therefore, we have good resolution to identify sensitive interactions, but not necessarily resistance. Third, unknown off-target effects can be difficult to estimate. We partly addressed this issue with deep learning predicted drug target which could be used for drug target discovery. Finally, in our analysis we had less than 50 samples for some tissues and used only 102 protein targets for the interaction matrix. Having more cell lines and more drugs should lead to improved predictability and more findings.

Multitask learning framework can handle very diverse prediction settings (**Supplementary Fig. S1**), and can be a useful tool for the advance of precision medicine. Depending on the availability of the data and objectives, it allows us to find genomically defined patients for existing drugs and ideal drugs for existing patients, as well as giving existing drugs to existing patients and test new drugs on new patients. Although our results are based on cell lines and hence unlikely to be directly suited for predicting clinical outcome, they can still be used for exploring mechanism of action of drugs and their contribution to the overall outcome.

We used PROGENy pathway activities derived from gene expression, as they provide a rich characterization of the cellular status. We also explored how mutations (SNP) and copy number variations (CNV) interact with drug targets (**Supplementary Fig. S6**), but the prediction performance (quality control) using SNP/CNV is generally lower than using PROGENy and that not all SNP/CNV are present in every tissue (**Supplementary Table S4, Methods**). For instance BCR-ABL mutation appears only for leukemia tissue (**Supplementary Fig. S6**), which makes this biomarker difficult to generalize to other cancer types. Prediction performance and availability of the features across tissues were the main reasons we did not focus on SNP/CNV in this paper.

Exploring the interactions between drug targets and signaling pathways can provide novel in-depth view of cellular mechanism and drug mode of action, which will ultimately rationalize tissue specific therapies. Feature interaction analysis based on deep learning predicted protein targets could potentially help target discovery. Previous studies showed the importance of network analysis to find new drug targets^24^. Zaman *et al.^25^* reported subtype-specific drug targets, which supports our result regarding the diversity of target - pathway interactions across different tissues. In cross tissue analysis (**Figure 4**), the triplet pathway/target/tissue allow drug repositioning and patient stratification strategies. It highlights cases of interaction that can provide useful biomarkers on one cancer type but potentially provide the inverse stratification for another cancer type, thus leading to treating the wrong patients. Knowing the variation of those interactions across tissues may be informative for drug repurposing, drug combination design and patient stratification.

## Methods

### Macau: Algorithm

Macau trains a Bayesian model for collaborative filtering by also incorporating side information on rows and/or columns to improve the accuracy of the predictions (**Figure 1a**). Drug response matrix (**IC50**) can be predicted using side information from both drugs and cell lines. We use protein target as drug side information and transcriptomics/pathway as cell line side information. Each side information matrix is then transformed into a matrix of L latent dimension by a link matrix. Drug response is then computed by a matrix multiplication of the 2 latent matrices. Macau employs Gibbs sampling to sample both the latent vectors and the link matrix, which connects the side information to the latent vectors. It supports high-dimensional side information (e.g. millions of features) by using conjugate gradient based noise injection sampler. For more information, see Supp methods.

### Feature interaction

#### Concept

We would like to know the interactions between the features of the drugs and the features of the cell lines. In our analysis, we used protein target to describe the drugs and gene expression/PROGENy pathways to describe the cell lines. Let IC50 be the matrix of drug response, D be the latent matrix of the drugs and C be the latent matrix of the cell lines (**Figure 1a**):

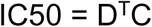

if side information (feature) are available on both sides:

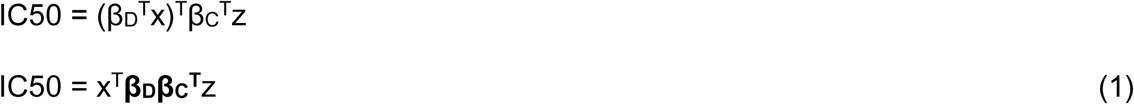

The matrix **β_d_β_c_^t^** is the interaction term or the interaction matrix through which the 2 feature sets interact in order to produce the response variable IC50 (**Figure 1b**). We generated the interaction matrix between features of the drugs and features of the cell lines by multiplying the 2 link matrices **β_d_** and **β_c_** and averaging across 600 MCMC samplings. We used setting 3 (**Supplementary Fig. S1c, Supplementary Table S1**) to compute the interaction matrix for the feature interaction analysis. This setting allows the use of the whole dataset, without cross validation. MCMC sampling is also less prone to overfitting than optimization methods.

Each drug response observation can be written as a linear combination of all the possible interactions between the protein targets and the pathways’ activity, across all latent dimensions.

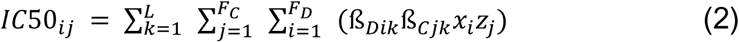

L: number of latent dimensions

F_d_: length of drug feature (number of protein targets)

F_c_: length of cell line feature (number of PROGENy pathways)

x: drug feature (protein target)

z: cell line feature (PROGENy pathway activity)

**β_d_**: link matrix which projects drug feature X_d_ into latent matrix D

**β_c_**: link matrix which projects cell line feature X_c_ into latent matrix C

#### Interpretation

The interaction matrix **β_d_β_c_^t^** has as dimension the number of protein targets multiplied by the number of pathways. We then multiply the matrix by −1 so that the interpretation would be:

In case of a positive value in the matrix, the association of the corresponding protein target and the corresponding pathway confers sensitivity upon drug treatment. If the value is negative, it would be resistance.

Another way to say it would be: If the value is positive, activation of this specific pathway confers sensitivity to any drug targeting this specific protein.

### Quality control of the interaction matrix

#### Significance of the method by cross validation

In order to assess the quality of the interaction matrix between drug targets and cell lines features, we used setting 4 i.e Predicting new drugs’ responses on new cell lines (**Supplementary Fig. S1d**) for each tissue type (**Supplementary Table S4**), since setting 4 describes the generalization to new cases, which is what we want to obtain with the interaction analysis. This does not give us a p-value for each value of the interaction matrix, but rather gives an overall quality of the model (pearson correlation of observed versus predicted IC50) for a given tissue and a pair of feature type. The performance across tissues are ranging from 0.33 for liver to 0.45 for skin. We consider a performance of 0.3 as a valid model. Setting 4’s double cross validation is the gold standard method for significance evaluation of feature interaction analysis. It is essential to perform this analysis first before considering the generated insights or looking into the significance of each value.

#### Significance of the result by a permutation based approach

We generated random permutations of the pathway activity matrix 1000 times, where we shuffled the PROGENy scores for each cell line independently. We did not randomise the drug target as we can lose the information that two drugs are targeting the same protein, which could be crucial in setting 4 (predicting new drug on new cell line). We then derived an empirical null distribution for each value of the interaction matrix. If the value is positive, we define the p value as the number of cases in the null distribution greater than the value of interest divided by 1000.

If the value is negative, we define the p value as the number of cases in the null distribution smaller than the value of interest divided by 1000. We then corrected for multiple non independent tests using the Benjamini-Hochberg Yekutieli procedure. We chose 20% as threshold of significant q value.

### Parameter setting

When predicting drug response on new cell lines (Supplementary Fig. S1a), we set the number of latent dimension L to 10 if we only use cell line feature. In case of adding drug feature, we set L to 30. Smaller L could lead to overcrowded latent space and decrease of performance. In MCMC sampling, we chose a burn in of 400 samples, then we collected 600 samples. At each of those collected samples, we made the prediction and averaged across all 600 samples. In quality control of both sides of features, we used setting 4 (Supplementary Fig. S1d) and 2 simultaneous 10 fold cross validation and 30 latent dimensions. In feature interaction analysis, we used setting 3 (Supplementary Fig. S1c), predicting existing drug for existing cell line) with 30 latent dimensions.

### PROGENy

PROGENy^12^ is a data driven dimension reduction method for gene expression data. It reduces high dimensional gene expression into a small number of pathway activity scores by a matrix multiplication with a weight matrix. PROGENy leverages hundreds of perturbation experiments. For each experiment, we assign a manually curated pathway activation status. The chosen experiments have been treated by a perturbation agent which activates or inhibits one of the PROGENy pathways.

We compute the gene expression z-scores of the Microarray_perturbed_ - Microarray_control_. Then, we fit a multiple linear model of the z-scores in function of the pathway status (**Supplementary Fig. S7a**). The z-scores representing the change in gene expression, we aim at determining the role of the pathway activation statuses in this change.

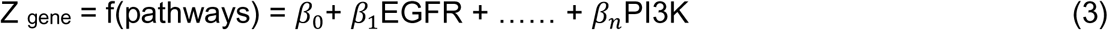

We obtain a pathway weight matrix from the fitted model. And for each pathway we select the 100 smallest p-values and keep those genes while setting the other genes’ weights to zeros (**Supplementary Fig. S7b**).

For new gene expression data where we would like to know the pathway information, we compute the pathway scores by multiplying the gene expression matrix with the pathway weight matrix. If we take the example of EGFR, the pathway activity of EGFR on sample 1 (s1) is defined as:

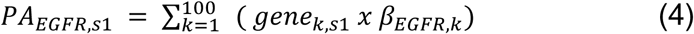

The pathway activity is defined as the product of a gene’ expression by the contribution of a pathway’s activation to the change in expression of this gene. From this formula, the higher the gene expression, the higher the pathway activity. Similarly, the higher the contribution of EGFR’s activation to the change of gene expression, the higher the pathway activity.

The result is the pathway scores matrix with new experiments on the rows and pathways on the columns (**Supplementary Fig. S7c**). In practice, for any transcriptomics dataset, we can determine which pathway is up regulated or down regulated for a certain cell line relative to other cell lines. In this paper, we are using the pathway scores as features to predict drug response on cell lines. Therefore, PROGENy is used as a data driven dimension reduction method.

We compared PROGENy’s drug response prediction performance with pathway scores calculated from other approaches, such as Biocarta, Gatza et al, Gene Ontology, Reactome, PARADIGM, Pathifier, and SPIA. We obtained the signature score for those methods as described in Schubert *et al*. We used those features as input for drug response prediction on GDSC dataset. For the 11 essential cancer pathways we used in our analysis, PROGENy performed better than all other methods (**Supplementary Fig. S9**). It performed significantly better than the second best pathway score Biocarta (p=0.00038).

### Data

GDSC data were downloaded from: http://www.cancerrxgene.org/

Drug IC50 version 17a
Basal gene expression 12/06/2013 version 2
Drug target version March 2017

CTRPv2 data were downloaded in 2016 from: https://portals.broadinstitute.org/ctrp/

Seashore-Ludlow *et al*., 2015

### Availability and Implementation

The source code of the method is available at https://github.com/saezlab/Macau_project_1

## Acknowledgements

The work received funding through the JRC for Computational Biomedicine which was partially funded by Bayer AG. We would like to thank Bence Szalai, Satya Swarup Samal, Vigneshwari Subramanian and Damien Arnol for their suggestions and ideas in improving the manuscript.

## AUTHORS CONTRIBUTIONS

MY designed research, performed all analyses, and wrote the manuscript. JS developed Macau algorithm, CCL performed the target prediction, PZ wrote supplementary method, GJW wrote supplementary method, YM supervised the development of Macau algorithm, JSR supervised the project and contributed to writing the manuscript.

